# Lung endothelium instructs dormancy of susceptible metastatic tumour cells

**DOI:** 10.1101/2022.06.18.496680

**Authors:** Moritz Jakab, Ki Hong Lee, Alexey Uvarovskii, Svetlana Ovchinnikova, Shubhada R Kulkarni, Till Rostalski, Simon Anders, Hellmut G Augustin

## Abstract

During metastasis, cancer cells hijack blood vessels and travel via the circulation to colonize distant sites^1,2^. Due to the rarity of these events, the immediate cell fate decisions of arrested circulating tumour cells (aCTC) are poorly understood and the role of the endothelium, as the interface of dissemination, remains elusive^3,4^. Here, we developed a novel strategy to specifically enrich for aCTC subpopulations capturing all cell states of the extravasation process and, in combination with single cell RNA-sequencing, provide a first blueprint of the transcriptional basis of early aCTC decisions. Upon their arrest at the metastatic site, tumour cells either started proliferating intravascularly or extravasated and preferably reached a state of quiescence. Endothelial-derived angiocrine Wnt factors were found to drive this bifurcation by inducing a mesenchymal-like phenotype in aCTCs instructing them to follow the extravasation-dormancy branch. Surprisingly, homogenous tumour cell pools showed an unexpected baseline heterogeneity in Wnt signalling activity and epithelial-to-mesenchyme-transition (EMT) states. This heterogeneity was established at the epigenetic level and served as the driving force of aCTC behaviour. Hypomethylation enabled high baseline Wnt and EMT activity in tumour cells leading them to preferably follow the extravasation-dormancy route, whereas methylated tumour cells had low activity and proliferated intravascularly. The data identify the pre-determined methylation status of disseminated tumour cells as a key regulator of aCTC behaviour in the metastatic niche. While metastatic niche-derived factors per default instruct the acquisition of quiescence, aCTCs unwind a default proliferation program and only deviate from it if hypomethylation in key gene families renders them responsive towards the microenvironment.

## Main

Tumour cell (TC) dormancy poses a major hurdle for the treatment of cancer^1-3^. During metastasis, dormant tumour cells (DTC) reside in close proximity to blood vessels and acquire a stem-like phenotype^4^. Yet, metastasizing TCs show a remarkable heterogeneity in their genetic and molecular makeup, which can be attributed to why certain cancer cells reach a state of tumour dormancy, whereas others outgrow immediately to form macro-metastases^5-8^. This differential behaviour is not solely driven by TC intrinsic properties, but also influenced by the niche, as certain niches in principle favour TC proliferation, whereas others are primarily tumour-suppressive^9-11^. This argues for a scenario in which the induction of dormancy depends on cell-intrinsic properties and matching microenvironmental factors. This is also corroborated by the finding that DTCs, once committed to their fate, require dramatic events to be awakened^12,13^. We therefore hypothesized that TC behaviour is primed during the initial arrival of CTCs at the metastatic niche and that the vascular endothelium, as the interface of dissemination, is a crucial fate instructor.

### Wnt and EMT pathways are drivers of tumour extravasation and dormancy

To test our hypothesis, we developed a novel experimental model to temporally assess TC and endothelial cell (EC) interactions *in vivo* at single cell resolution for the first time. Wildtype Balb/c mice were intravenously injected with GFP-expressing 4T1 breast cancer cells (4T1-GFP). Lung-seeded TCs and corresponding total lung ECs were isolated at day 0 (baseline), day 1.5 (peak phase of extravasation) and day 3.5 (induction of proliferation) and purified by fluorescence-activated cell sorting (FACS) (**Fig. 1a**). Plate-based single cell RNA-sequencing (scRNAseq) was used to analyse the transcriptional signatures of TC-EC interactions during metastatic colonization. To discriminate extravascular from intravascular TCs, mice were intravenously injected at day 1.5 with fluorescently labelled anti-H2-Kd antibody labelling all body cells, including syngeneic 4T1-GFP cells that were exposed to the circulation. We further discriminated proliferative from quiescent TCs at day 3.5 based on the dilution of CellTracer dye and considered TCs exhibiting dye retention as dormant (**Fig. 1a**). For each TC subpopulation and matched ECs, equal numbers were sorted from at least three biological replicates. The cells in the dataset showed homogenous distribution for counts and detected genes and the replicates interlaced well in a Uniform Manifold Approximation and Projection (UMAP), demonstrating the robustness of the experimental approach. Moreover, substructures in the UMAP were found to be specifically enriched for cells from the respective FACS gates, suggesting that the gating strategy was suitable to enrich rare TC subpopulations, even though it did not yield high purity. Correspondingly, *H2-K1* expression did not differ between the intravascular and extravascular fraction, highlighting that differences in staining intensity reflected exposure to circulation and not gene regulation.

**Fig. 1:**
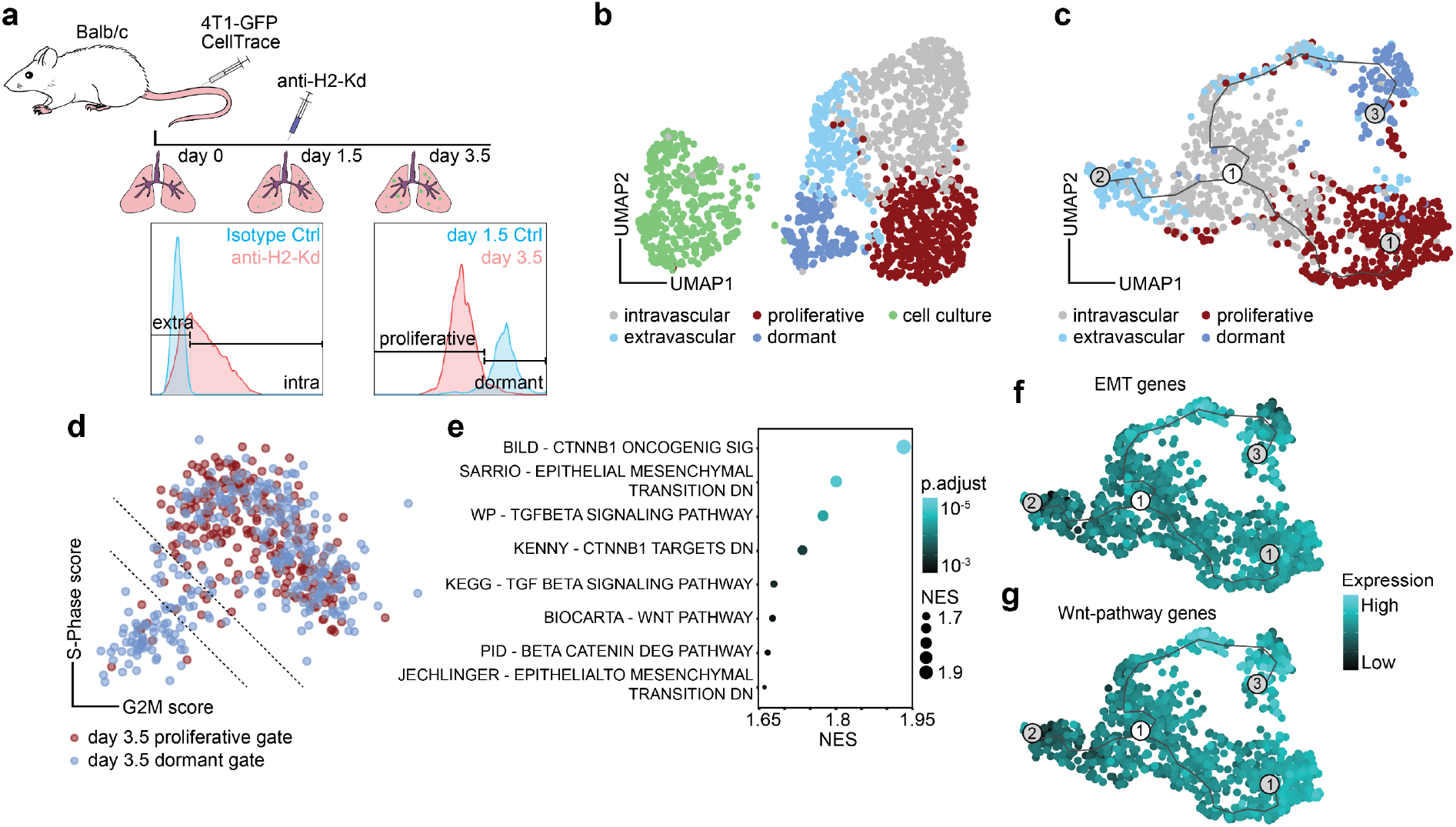
Extravasating and dormant tumour cells are defined by a Wnt and EMT signature. **a**, Schematic of the experimental design. Recipient Balb/c mice received two injections of 1×10^6^ 4T1-GFP cells stained with CellTracer dye into the tail vein. At day 1.5 post-injection, mice were injected intravenously with 5µg of fluorescently labelled anti-H2-Kd antibody. Tumour cells (TC) and endothelial cells (EC) were sampled at day 0 (uninjected baseline), day 1.5 and day 3.5. TCs were discriminated based on extravasation status on day 1.5 and based on proliferation status on day 3.5. **b**, Uniform manifold approximation and projection (UMAP) of total TC dataset. Shared nearest neighbour (SNN-) based clustering resolves transcriptomes of TCs into 5 clusters. **c**, Trajectory analysis of extracted TCs reveals three transition branches. **d**, Scatter plot of S-Phase and G2M gene expression scores for individual cells extracted on day 3.5 and coloured by respective FACS gates. Dotted line indicates thresholds of cells with score sums < −1 (lower line) and score sums < 0 (upper line). **e**, Gene set enrichment analysis (GSEA) of genes upregulated in bona fide dormant tumour cells ranked by fold change. NES, normalised enrichment score. **f**, Summed expression of 384 genes upregulated during epithelial-to-mesenchyme transition (EMT) and **g**, 146 genes associated with Wnt pathway plotted on the trajectory graph from **c**.

Clustering of the combined TC dataset identified a total of five clusters that reflected the respective FACS gates (**Fig. 1b)**. Trajectory analysis to reconstruct the colonization process with TCs of the intravascular cluster set as starting point identified three main trajectories transitioning 1) from intravascular to proliferative cells, 2) to a subset of extravascular cells, and 3) through a subset of extravascular cells to dormant cells (**Fig. 1c**). These findings indicated that TC extravasation was a pre-requisite for tumour dormancy, but dispensable for TC proliferation, which molecularly defines earlier microscopy-based concepts^14,15^. Based on this finding, we next compared proliferative vs. dormant cells and scored each TC from the day 3.5 dataset for the expression of G2M-phase and S-phase genes. Only TCs that showed dye retention and were not in cycle were considered as bona fide dormant to exclude cells that had proliferated but dropped out of cycle (**Fig. 1d**). After regression of cell cycle-associated genes, differential gene expression analysis (DGEA) was performed with subsequent gene set enrichment analysis (GSEA) (**Fig. 1e**). In line with previous reports^16,17^, transforming growth factor beta (TGFβ) signalling and epithelial-to-mesenchyme transition (EMT) gene sets were enriched in DTCs. Surprisingly, beta-Catenin-mediated canonical Wnt signalling was identified as one of the most significantly enriched pathways (**Fig. 1e**). Wnt ligands have extensively been characterized as protumorigenic growth factors^18^, promoting proliferation in primary tumours and metastases, as well as CTC survival^19-22^. Unexpectedly, expression of Wnt pathway and EMT-associated genes was enriched alongside the extravascular-dormancy trajectory (**Fig. 1f, g**), supporting the hypothesis that niche-derived Wnt ligands may drive tumour dormancy. In conclusion, these data provide a first transcriptional blueprint of aCTC fate decisions in the metastatic lung.

### The lung endothelium displays a bimodal response towards arriving CTCs

It was previously established that the endothelium serves as a systemic amplifier of primary tumour-derived signals^23,24^. Here, clustering of lung ECs revealed the emergence of cycling ECs as well as a shift of general capillary ECs (gCap)^25,26^ towards an activated phenotype as a response towards arriving TCs. To determine the basis of gCap activation, DGEA on filtered capillary EC pseudo-bulks was performed. For this, large vessel EC were manually annotated and removed from the dataset based on congruent marker gene expression. Capillary ECs displayed an immediate response pattern with genes being mostly regulated at day 1.5. These immediate-response genes involved immune modulatory and cell cycle genes and were regulated in a bimodal-manner with systemic upregulation of secreted EC factors (angiokines) and focal enrichment of biosynthesis genes in activated gCap. This led us to conclude that the lung endothelium, while exerting important immune regulatory functions at the systems level, also served as a local producer of biomass thereby generating a conducive micro-niche.

### Niche-derived angiocrine Wnt ligands are instructors of tumour dormancy

Next, we analysed the consequences of the enriched Wnt signalling in DTCs. For this purpose, 4T1-GFP breast cancer cells were treated *in vitro* for 2 weeks with canonical Wnt pathway agonists prior to injection in a gain-of-function (G-O-F) approach. Conversely, mice were treated with a Porcupine inhibitor (LGK974) to create a Wnt-deficient environment (**Fig. 2a**). As expected, Wnt G-O-F programmed TCs to follow the extravasation-dormancy route, which resulted in enhanced extravasation and higher incidence of dormancy with overall reduced short-term metastatic burden (**Fig 2b, c)**. In contrast, Wnt depletion enhanced metastatic outgrowth, but did not affect extravasation (**Fig 2b, c**). As modulating Wnt signalling was sufficient to alter TC behaviour *in vivo*, we probed for sources of Wnt ligands in the metastatic niche. The endothelium was found to robustly express Wnt ligands across the experimental timeline. Yet, their expression was not changed (**Fig. 2d)**. Nevertheless, depleting Wnt ligands specifically from the vascular niche by EC-specific knockout of the Wnt cargo receptor *Wntless* (*Wls*) led to a significantly increased short-term metastatic burden, thereby phenocopying the systemic pharmacological inhibition. This was observed for E0771-GFP breast cancer cells, but also for B16F10 melanoma cells (**Fig. 2e**), highlighting the endothelium as a major source of dormancy-inducing Wnt ligands. The Wnt dependency of metastatic dormancy acquisition could also be observed in a clinically relevant spontaneous metastasis model involving surgical removal of the primary tumour (**Fig. 2f**). Specifically, systemic treatment with LGK974 did not affect primary tumour growth or size, mouse body weight or the tumour vasculature (**Fig. 2g)**, but resulted in a significantly increased metastatic burden (**Fig. 2h**). Collectively, these data establish that angiocrine Wnt ligands are crucial instructors of extravasated TC dormancy.

**Fig. 2:**
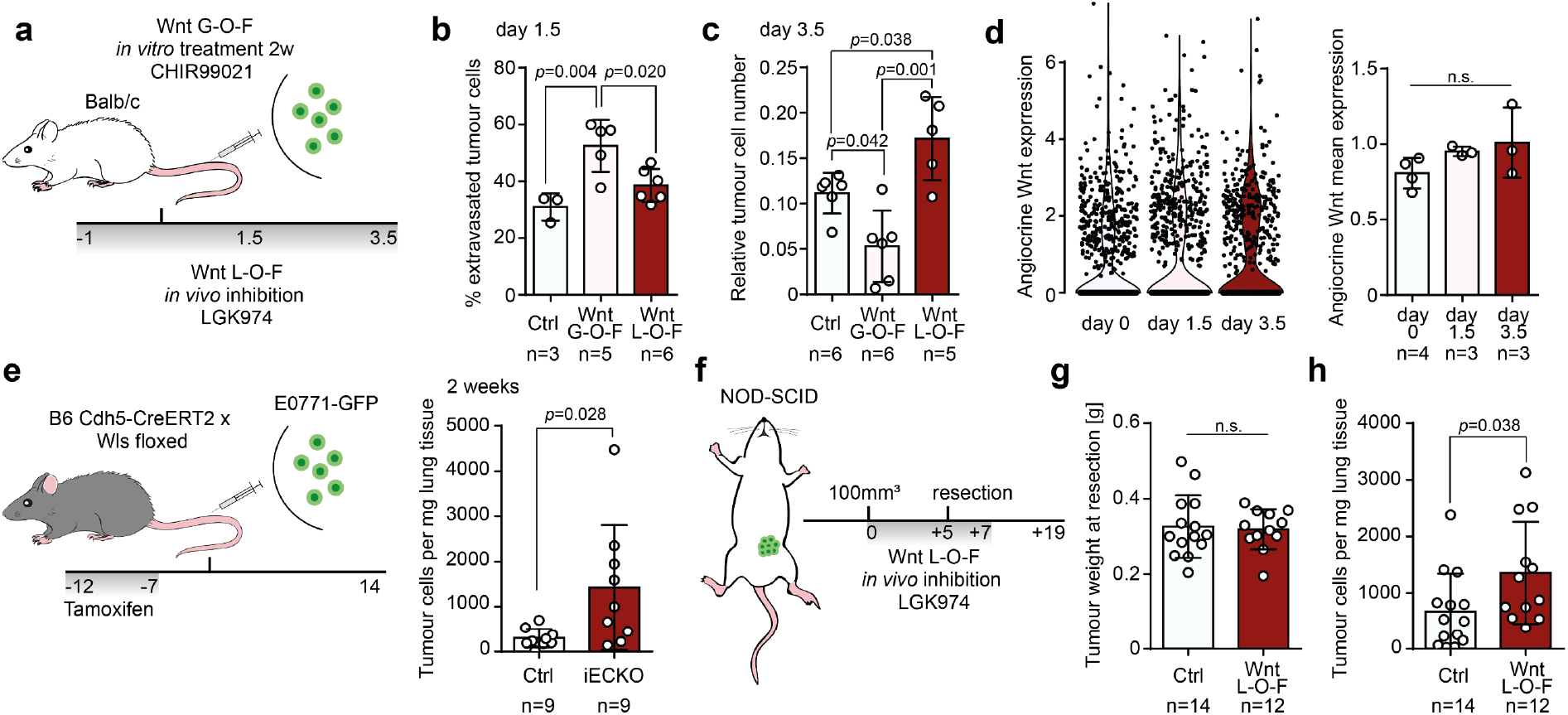
Lung endothelial cells are a major source of dormancy-inducing Wnt ligands. **a**, Schematic of the experimental gain-of-function (G-O-F) and loss-of-function (L-O-F) strategy. Grey bar indicates timespan (in days) of daily treatment with LGK974. **b**, Percentage of extravasated TCs for control treatment, G-O-F or L-O-F approach 1.5 days post-injection. Error bars s.d., *p* values by one-way ANOVA with Tukey post-test are shown. **c**, Quantification of relative TC number normalised to EC abundance in lungs 3.5 days post-injection for control treatment, G-O-F or L-O-F approach. Error bars s.d., *p* values by one-way ANOVA with Tukey post-test are shown. **d**, Summed expression of Wnt ligands per timepoint in individual ECs (left panel) or as mean expression for pseudo-bulks of biological replicates. Error bars s.d., *p* values were calculated by one-way ANOVA with Tukey post-test. n.s., not significant. **e**, Schematic of experiment. Gene recombination was induced by tamoxifen administration. 2×10^5^ E0771-GFP cells were injected into the tail vein of EC-specific knockout (iECKO) and control animals. Grey bar indicates timespan (in days) of daily tamoxifen treatment (left panel). Total number of TCs per milligram lung tissue of control and iECKO mice two weeks post-injection of E0771-GFP. Error bars s.d., *p* value by two-tailed *t*-test is shown. **f**, Schematic of the experiment. 1×10^6^ 4T1-GFP cells were implanted into the mammary fat pad of NOD-SCID mice. Once tumours reached a size of 100mm^3^, mice were treated with LGK974 for five days until tumour-resection. After resection, mice were treated for an additional two days and left to develop metastases. Grey bar indicates timespan (in days) of daily treatment with LGK974. **g**, Weights of resected primary tumours. Error bars s.d., *p* value was calculated by two-tailed t-test. n.s., not significant. **h**, Total number of TCs per milligram lung tissue of control and LGK974 treated mice two weeks post-resection. Error bars s.d., *p* value by two-tailed t-test is shown.

### Metastatic tumour cell behaviour is not regulated at the receptor-ligand level or by niche occupancy

As the expression of angiocrine Wnt ligands was not changed, we reasoned that Wnt signalling activity differences may result from distinct TC receptor repertoires. Surprisingly, prediction of EC-TC interactions based on DGEA of TC pseudo-bulks (intravascular vs extravascular and proliferative vs. dormant) using CellPhoneDB^26^ did not reveal Wnt pathway components. Repeating the analysis specifically with TC-expressed Wnt receptors led to the identification of five (co)-receptors that were differentially expressed for either the intravascular vs extravascular or the dormant vs proliferative comparison, but not for both. Moreover, receptor expression was not enriched in the intravascular-proliferative or the extravasation-dormancy branch of the trajectory-analysis, suggesting that the observed differences in Wnt signalling activity were not established at the receptor-ligand level.

As the lung endothelium harbours two distinct vascular beds that are defined by less penetrable gCaps and more permissive aerocytes (aCap)^23, 24^ (**Fig. 3a**), we tested whether distinct vascular niche occupancy could drive the observed differential TC behaviour. Employing an *in vivo* niche-labelling system^28^, we specifically enriched for ECs interacting with mostly proliferative 4T1-GFP cells or quiescent D2.0R-GFP breast cancer cells that served as a proxy for dormant 4T1 cells. Labelled niche ECs, as well as matched unlabelled total ECs were FACS-purified and subjected to bulk RNAseq (**Fig. 3b, c**). Aerocyte and gCap-specific genes that showed robust and stable expression were used to deconvolute the cellular composition of the bulk samples (**Fig. 3d**). While a general bias towards the aCap signature could be observed for all tumour-bearing samples, no differences were detected between the dormant and proliferative niche or their unlabelled counterparts (**Fig. 3e**), indicating that DTCs and proliferative TCs occupy the same vascular niches.

**Fig. 3.**
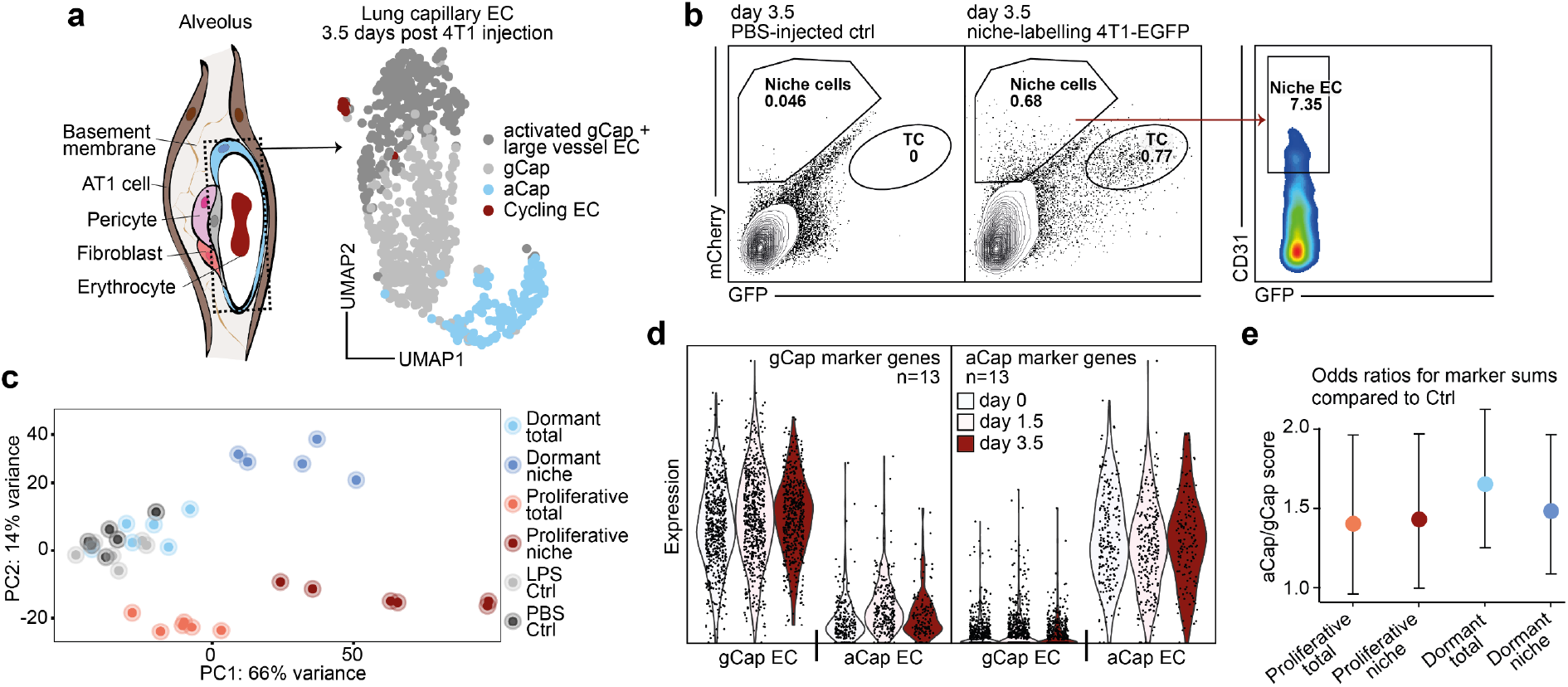
Dormant tumour cells do not occupy distinct vascular niches in the lung. **a**, Schematic of lung alveolus (left panel). Dotted box highlights ECs. UMAP of EC transcriptomes reflecting the composition of bulk EC samples 3.5 days post-injection of 4T1-GFP cells. **b**, Representative FACS gates for purifying labelled niche ECs 3.5 days post-injection of niche-labelling 4T1-GFP. **c**, Principal component analysis of samples included in the experiment. Total samples refer to unlabelled CD31+ ECs, niche samples refer to labelled CD31+ ECs, dormant samples refer to injections of niche-labelling D2.0R-GFP, proliferative samples refer to injections of niche-labelling 4T1-GFP, LPS controls were injected intraperitoneally with LPS 24 hours prior to euthanasia, PBS controls were injected intravenously with PBS 3.5 days prior to euthanasia. **d**, Summed expression of gCap and aCap marker genes in individual ECs split by timepoints and by EC identity. **e**, Log2 fold changes of aCap marker genes to gCap marker genes odds ratios normalised to PBS-injected control samples. Error bars indicate 95% confidence interval.

Interestingly, proliferative TCs, in contrast to DTCs, induced the production of extracellular matrix (ECM), immune response genes and proliferation in ECs. While matrix-remodelling processes were specific to the niche, proliferative and pro-inflammatory programs were part of a systemic response. We sought to deduce a marker gene set from the bulk comparisons that was specific for ECs extracted from the proliferative niche and tested all conditions against each other. The resulting gene panel was used to predict tumour-interacting ECs in the scRNAseq data. Predicted tumour-interacting ECs co-localized with the previously identified biosynthetic ECs in the UMAP, confirming the hypothesis that the endothelium elicited a bimodal response. To test the generality of the gene panel, a publicly available scRNAseq dataset^29^ was utilized and a similar enrichment was found specifically for primary lung tumour ECs compared to non-tumorous matched samples.

### Heterogenous methylation states pre-determine tumour cell behaviour

As DTC fates were established independently of differential exogenous factors, we next probed for TC-intrinsic properties. Surprisingly, cultured TCs exhibited a heterogeneous but correlating baseline expression of EMT-and Wnt pathway-associated genes (**Fig. 4a, b**). Such state differences were also identified in freshly isolated CTC of breast cancer patients^30,31^, indicating baseline tumour cell-intrinsic differences. Interestingly, overnight pulse treatment with Wnt agonists failed to programme the extravasation-dormancy shift observed for long-term treatments (**Fig. 4c)** and the expression profile of key EMT-associated transcription factors was not markedly changed between pulse-treated and reprogrammed cells. This led us to hypothesize that TC were restricted in their responsiveness towards niche-derived factors by an epigenetic barrier. In agreement, pulse-treatment of 4T1-GFP with de-methylating agent (decitabine) enabled TCs to preferably follow the extravasation-dormancy route (**Fig. 4d**). This was not driven by cellular fitness or changes in homing capacity. Similar to the Wnt G-O-F, hypomethylation did not induce EMT *in vitro* and dormancy-induction was still dependent on Wnt (**Fig. 4e**). Notably, pulse-treating hypomethylated cells with Wnt agonist prior to injection did not alter the *in vivo* phenotype, indicating that niche-derived signalling was saturated and sufficient. Moreover, none of the *in vitro* treatments affected the proliferation rate of proliferation-committed TCs *in vivo*, clearly showing that differences in short-term metastatic burden were a consequence of dormancy-induction.

**Fig. 4.**
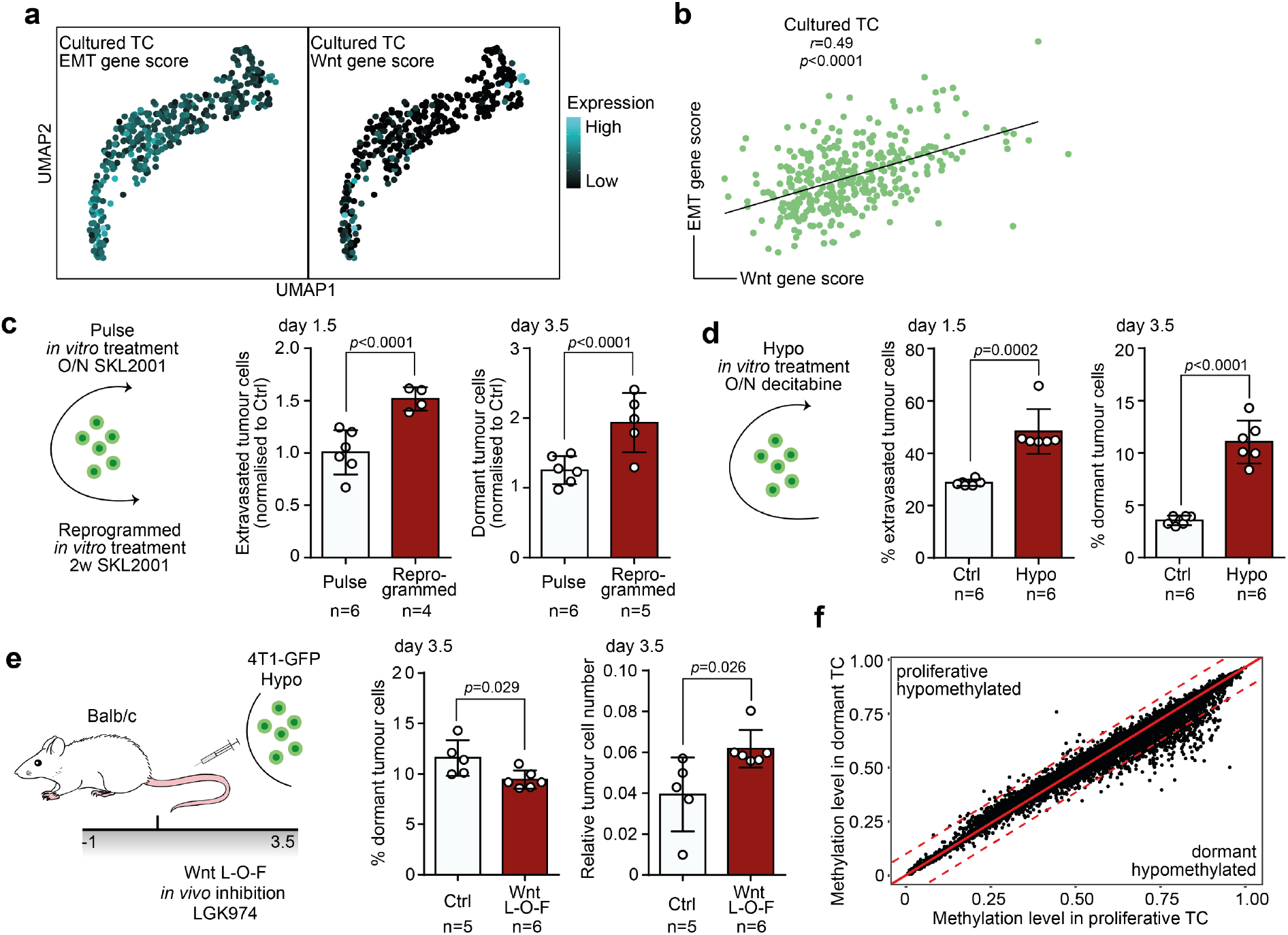
Tumour cell behaviour in the metastatic niche is pre-determined by methylation state. **a**, Gene scores of EMT (left panel) and Wnt pathway-associated genes (right panel) in cultured TCs visualised in a UMAP and **b**, correlation of the gene scores. *P* value and *r* value by Pearson correlation are shown. **c**, Schematic of experiment (left panel). 4T1-GFP cells were either treated overnight with Wnt agonist (pulse) or for two weeks (reprogrammed) prior to injection. Relative fraction of extravasated TCs normalised to the respective control 1.5 days post-injection (mid panel) and relative fraction of dormant TCs normalised to the respective control 3.5 days post-injection (right panel). Error bars, s.d., *p* values by two-way ANOVA with Sidak post-test are shown. **d**, Schematic of experiment (left panel). 4T1-GFP cells were treated overnight with de-methylating agent decitabine (hypo). Percentage of extravasated TCs for control and hypomethylation treatment 1.5 days post-injection (mid panel) and percentage of dormant TCs 3.5 days post-injection (right panel). Error bars, s.d., *p* values by two-tailed t-test are shown. **e**, Schematic of experimental L-O-F approach (left panel). 4T1-GFP cells were treated overnight with decitabine. Grey bar indicates timespan (in days) of daily treatment with LGK974. Percentage of dormant TCs 3.5 days post-injection for control and LGK974 treated animals (mid panel) and relative tumour cell number normalised to EC abundance in lungs (right panel). Error bars, s.d., *p* values by two-tailed t-test are shown. **f**, Scatter plot of methylation level (fraction of methylated CpG islands) for gene bodies in dormant TCs and proliferative TCs. Red line indicates no differences in methylation, dotted red lines indicate thresholds for >10% differences in methylation.

We then assessed the methylation state of dormant and proliferative TCs by whole genome bisulfite-sequencing. Global methylation levels were not changed in DTCs, but promoter sequences and gene bodies displayed considerable hypomethylation, while enhancer sequences were unaffected (**Fig. 4f)**. Remarkably, hypomethylation mainly occurred in genes and promoters that were epigenetically sealed in proliferative TCs (>70% methylation). We computed the overlap of genes with >10% hypomethylation in DTC and gene ontology terms from the mouse signature database (MSigDB)^32^ and found transcription factor binding and cell fate processes as top hits. This confirmed the hypothesis that hypomethylation underlay cellular plasticity and was the driving force of TC responsiveness towards niche-derived dormancy-inducing factors.

## Discussion

Tumour cell pre-determination is an emerging concept^33-35^. Here, we identified the epigenetic pre-coding of disseminated TC behaviour in the metastatic niche. DTC were characterized by hypomethylation in promoter sequences and gene bodies, whereas proliferative TCs were epigenetically sealed. We envision that the plastic-dormant and the sealed-proliferative state form a dynamic equilibrium. Long-term treatment with a Wnt agonist would direct the equilibrium towards the plastic-dormant state, without affecting the cell-state itself. This is supported by reports of similar phenotype transitions caused by long-term *in vitro* treatments or targeted genetic manipulation of signalling pathways^36,37^. However, such state-transitions were also reported to occur spontaneously and were found to be a pre-requisite for metastatic outgrowth^38^. TC hypomethylation was reflected at the transcriptomic level by an elevated baseline expression of EMT and Wnt pathway-associated genes. A similar heterogeneous expression was found in freshly isolated CTCs from breast cancer patients and could be linked directly to their metastatic potential^30,39^. In this context, the primary tumour could be viewed as a heterogenous amplifier in which high selective pressure forces the acquisition of distinct TC states. Recent lineage tracing experiments highlighted this phenomenon and revealed hybrid EMT TC states as the underlying principle of metastatic dissemination^5-8,35^. While EMT was needed for migration and intravasation, too much of it limited metastatic outgrowth. Interestingly, hybrid EMT states were not discrete but formed a continuum that correlated with the metastatic outcome^6-8^. Similar gradual states could be observed in cultured TCs and could be a direct consequence of epigenetic plasticity. Plastic cells would show high EMT and follow the extravasation-dormancy route, whereas sealed TCs would form macro-metastases. Probing the epigenetic and transcriptomic state of CTCs could therefore serve as predictive tool to assess the likelihood of metastatic relapse in patients.

Besides cell-intrinsic properties, disseminated TC phenotypes are established as a consequence of instructive niche-derived factors. Here, we identified endothelium-derived angiocrine Wnt signalling as a prototypic example of such dependency. However, other factors and other cellular sources have been identified previously and are most likely to act synergistically^9, 16, 17, 40-44^. Most surprisingly, homeostatic angiocrine Wnt signalling was found to be sufficient to drive dormancy-induction, suggesting a default tumour-suppressive lung niche. Similar default programs could occur in other organs in which ECs comprise a major Wnt source and were reported previously in different contexts^9,11^. Moreover, primary tumour-instructed remodeling of the niche could change the default state^45-49^. Additionally, the data strongly suggest that metastatic TCs actively inflicted a niche-EC gene program that resembled primary tumour EC signatures^29,50^ and that could fuel TC proliferation by altering the biophysical properties of the micro-niche.

Collectively, these data provide a first insight into the establishment of arrested CTC fates. We show that susceptible epigenetic states render TCs responsive towards niche-derived default factors, thus, opening the opportunity to probe for TC-niche interdependencies at the systems level.

## Acknowledgements

The authors would like to thank Dr. Kairbaan Hodivala-Dilke (Barts Cancer Institute) and Dr. Ilaria Malanchi (Francis Crick Institute), Dr. Jonathan Sleeman (University Medical Center Mannheim), and Dr. Robert Weinberg (Whitehead Institute) for providing cell lines and reagents. We thank Celine Rausch, Clara Mai and Carleen Spegg for technical assistance. We are grateful for the technical support of the Flow Cytometry Core Facility, the Single-Cell Open Lab, the Genomics and Proteomics Core Facility, the Omics IT and Data Management Core Facility, the Laboratory Animal Core Facility and the Light Microscopy Core Facility of the DKFZ. This work was supported by grants from the Deutsche Forschungsgemeinschaft (Collaborative Research Center CRC1366 ‘Vascular Control of Organ Function’ [project number 39404578; projects C5 to H. G. Augustin and Z3 to S. Anders], Collaborative Research Center CRC1324 ‘Wnt signaling’ [project number 331351713; project A2 to H. G. Augustin], the European Research Council Advanced Grant “AngioMature” [project 787181 to H. G. Augustin], the Deutsche Krebshilfe grant “AgedSoil” within the Excellence Program for Established Scientists [project 70114532 to H.G. Augustin] and the State of Baden-Württemberg Foundation special program “Angioformatics Single Cell Platform” [to H.G. Augustin].

## Data availability

All raw sequencing data, annotated and filtered count matrices generated in this study, will be made available to the reviewers upon request and publicly available upon final acceptance of the manuscript.

## Code availability

All code generated in this study will be made available to the reviewers upon request and publicly available upon final acceptance of the manuscript.

## Materials & Methods

### Animal studies

Female NOD-SCID and BALB/c mice were acquired from Janvier Labs. B6 *Cdh5*-CreERT2 x *Wls* floxed mice were published previously^51^ and bred in barrier animal facilities of the German Cancer Research Centre. All animal work was performed in accordance with German national guidelines on animal welfare and the regulations of the regional council Karlsruhe under permit numbers G-164/16, G-107/18, DKFZ305 and DKFZ370. Mice were housed in sterile cages, maintained in a temperature-controlled room and fed autoclaved water and food *ad libitum*. All animals were monitored daily for signs of disease and ear punches were used for genotyping the mice. Imported mice were allowed to acclimatize for a minimum of seven days before each experiment. For all experiments, 8-12 weeks old mice were used and euthanized via rapid cervical dislocation of the spinal cord at the experimental endpoint.

For experimental metastasis, tumour cells were resuspended in 200 µl PBS and injected into the tail vein of mice. For transcriptomic and epigenomic screening experiments (**Fig. 1a, Fig. 3a**), female Balb/c mice were injected twice with 1×10^6^ tumour cells (4T1-GFP, niche-labelling 4T1-GFP, niche-labelling D2.0R-GFP) with a 30 minute break between injections. For pharmacological treatment studies, female Balb/c mice were injected once with 1×10^6^ 4T1-GFP cells. For genetic knockout experiments, male and female B6 *Cdh5*-CreERT2 x *Wls* floxed mice were injected with 2×10^5^ E0771-GFP or B16F10, respectively. To stain intravascular cells, 5 µg of fluorescently labelled anti-H2-Kd antibody in 50 µl PBS was injected retro-orbitally into mice 2 min prior to euthanasia.

Pharmacological systemic depletion of WNT was achieved by daily oral gavage of 10 mg/kg body weight LGK974 resuspended in 0.5% methylcellulose (Sigma-Aldrich), 0.5% TWEEN 80 (Sigma-Aldrich) in PBS as described previously^52^.

EC-specific depletion of Wnt ligands was achieved using B6 *Cdh5*-CreERT2 x *Wls* floxed mice. Genetic recombination was initiated by intraperitoneal delivery of 2 mg tamoxifen (Sigma Aldrich) dissolved in 50 µl corn oil with 5% ethanol. Both Cre+ and Cre-littermates received 5 consecutive daily injections and were subjected to a one-week washout period before the start of the experiment.

To model spontaneous dissemination, 1×10^6^ 4T1-GFP cells in 100 µl PBS were injected into the inguinal mammary fat pad of female NOD-SCID mice. Tumour volumes were assessed by calliper measurements (tumour volume = ½ x length x width x width). Upon reaching tumour sizes of 100 mm^3^, mice were treated daily with LGK974 for 5 days with subsequent resection of the primary tumour. LGK974 treatment continued for 2 days post-resection and mice were left to develop metastases for 2 weeks.

To account for general inflammatory signatures in lung EC in the niche-labelling experiment (**Fig. 3b, c**), mice were injected with 1 mg/kg LPS (Sigma Aldrich) in 0.9% NaCl (Braun) intraperitonially 24 h prior to euthanasia.

After euthanizing the mice, lungs were collected in PBS, metastatic foci were counted (B16F10 experiments), lungs were imaged using a stereomicroscope (Leica) (E0771 experiments, primary tumour experiments) and processed for flow cytometry. Resected primary tumours were rinsed in PBS and fixed in formalin free Zn-buffer.

### Cell Culture

4T1-GFP, D2.0R and E0771-GFP cells were a gift from the laboratories of Dr. Robert Weinberg (Whitehead Institute, Cambridge, MA), Dr. Jonathan Sleeman (Heidelberg University, Mannheim, Germany) and Dr. Kairbaan Hodivala-Dilke (Barts Cancer Institute, London, England), respectively. B16F10 cells were purchased from ATCC. All cells were maintained at 37°C and 5% CO_2_ in high humidity and cultured in high glucose DMEM (Gibco) supplemented with 10% (vol/vol) FCS and 100 U/mL penicillin/streptomycin (Sigma Aldrich). D2.0R-GFP and 4T1-mCherry cells were generated by lentiviral transduction with TurboGFP and mCherry reporter, respectively. Niche-labelling cells were generated by lentiviral transduction of 4T1-GFP and D2.0R-GFP cells. Niche-labelling lentivirus was provided by the laboratory of Dr. Ilaria Malanchi (Francis Crick Institute, London, England). Cells were checked regularly for mycoplasma contamination by PCR and cell identity was confirmed by cell morphology. Cells were subcultured upon reaching 80-90% confluency by trypsin-EDTA (Sigma Aldrich) treatment.

For *in vitro* treatments, cell media were supplemented with 20 mM SKL2001 in DMSO (Selleckchem), 3 mM in DMSO CHIR99021 (Selleckchem) or 1 µM in PBS decitabine (Sigma Aldrich). Cells were treated overnight (∼17 h) for pulse treatments (SKL2001, CHIR99021, decitabine) or for 2 weeks with daily media changes for reprogramming (SKL2001, CHIR99021) with drugs or vehicle (solvent only).

### Isolation of lung cells

Isolated lungs were minced on ice using curved serrated scissors. The minced tissue was resuspended in DMEM supplemented with Liberase Thermolysin Medium enzyme mix (0.2 mg/ml, Roche) and DNAse I (0.2mg/ml, Sigma Aldrich) and incubated at 37°C first for 15 min and then again for 12 minutes. After each incubation, minced tissues were passed through 18G cannula syringes 30 times. After the second incubation, digested tissues were passed through 100 µm cell strainer to remove tissue debris and cell clumps. The following steps were performed on ice. The digestion reaction was quenched by adding FCS and samples were centrifuged at 4°C and 400 *g* for 4 min. Erythrocytes were lysed by resuspending the cell pellet in pre-chilled 1x ammonium chloride potassium (ACK) buffer. The reaction was quenched by adding ice-cold PBS, followed by centrifugation.

### Flow cytometry analysis and FACS sorting

Whole lung single cell suspensions were passed through a 40 µm cell strainer and preincubated with anti-mouse CD16/CD32 Fc block (1:100, Thermo Fisher Scientific) for 15 min in flow buffer (PBS supplemented with 5% (vol/vol) FCS) and, subsequently, with the appropriate antibody-mix for 20 min on ice.

For cell sorting and flow cytometry analysis, dead cells were excluded by staining with FxCycle™ Violet Stain (1:1000, Thermo Fisher Scientific) or Fixable Viability Dye eFluor™ 780 (1:1000, Thermo Fisher Scientific) according to the manufacturer’s instructions. All samples were gated on viable cells followed by exclusion of cell doublets and CD45+, LYVE1+, PDPN+ and TER119+ cells using the BD FACS Diva Software (BD Biosciences). For flow cytometry, samples were recorded on the BD LSR Fortessa or BD FACSCanto II cell analyser (both BD Biosciences) and flow data was analysed with FlowJo software (BD Biosciences, v10). Tumour cells frequencies were calculated either as percentage of sample-matched lung endothelial cells (relative tumour cell number) or as total tumour cell counts per mg lung tissue using CountBrightTM Absolute Counting Beads according to the manufacturer’s protocol. Cells were sorted using a BD bioscience Aria cell sorting platform (BD Biosciences) with 100 µm nozzle.

### Single cell RNA-sequencing

ScRNAseq on tumour cell subpopulations and endothelial cells was performed using a modified SMART-Seq2 protocol^53^. In brief, single cells were sorted directly into 96 well-plates containing 1 µl of lysis buffer per well, centrifuged and snap frozen in liquid nitrogen. For CD45-PDPN-LYVE-TER119-CD31+ EC, four plates (384 cells) were sorted for 3 biological replicates from day 1.5 and day 3.5 (total of 1152 cells per timepoint). Day 0 control samples were split and three plates (288 cells) were sorted for 2 biological replicates on day 1.5, as well as on day 3.5 (total of 1152 cells), respectively, to account for technical batch-effects. For TC, 1 plate of matched intravascular and extravascular fractions (each 96 cells) was sorted from 4 biological replicates (total of 384 cells per fraction) on day 1.5. Similarly, 1 plate of matched dormant and proliferative fractions was sorted from 4 biological replicates on day 3.5. Frozen plates were thawed on ice and oligo-dT-primer were annealed at 70°C for 3 min. 1.3 µl of reverse transcription mix with template-switching oligo was added to each well and isolated mRNA was transcribed to full-length cDNA. Full-length cDNA was then amplified by adding 2.4 µl of PCR mastermix to each well. Due to their low RNA-content, EC cDNA was amplified using 22 cycles, while TC cDNA was amplified with 18 cycles. Amplified cDNA was purified using AMPure XP beads (Beckman Coulter) and random wells were selected for quality control using 2100 Bioanalyzer (Agilent) and Qubit fluorometer (Thermo Fisher Scientific). DNA concentration for each well was measured using Quant-iT™ high sensitivity kit (Thermo Fisher Scientific) and concentrations were manually adjusted to 0.1 - 0.3 ng/µl. Tagmentation was performed using the Nextera XT DNA library preparation kit (Illumina) and a mosquito liquid handler (SPT Labtech). For this purpose, 1.2 µl of Nextera XT – TD buffer mix was added to each well of a 384 well plate with 0.4 µl of cDNA. For EC, 96 well plates from the individual biological replicates were pooled into one 384 well plate, whereas TC replicates were pooled according to sort gates. After tagmentation, customized i5 and i7 index primers were added, resulting in unique labelling of each well in the 384 well plate and tagmented cDNA was amplified using 14 PCR cycles. All uniquely labelled wells from each plate were pooled and multiplexed libraries were purified and quality controlled using TapeStation (Agilent) and Qubit fluorometer (Thermo Fisher Scientific), resulting in one multiplex per biological replicate for EC and one multiplex per FACS-sorted TC fraction. Multiplexes were then sequenced on individual lanes on a HiSeq2000 (Illumina) using V4 50 cycle single read kit generating approximately 500.000 reads per cell.

### Bulk RNA-sequencing of labelled niche EC

For the niche-labelling experiment 50.000 unlabelled lung EC and sample-matched total labelled EC were directly sorted as described above into RNase-free 1.5 ml microcentrifuge tubes containing 100 µl lysis buffer and immediately snap frozen on dry ice. For each condition 6 biological replicates were included. For control samples (LPS control and PBS control) 50.000 lung EC were sorted. Snap frozen RNA was extracted using Arcturus PicoPure RNA Isolation Kit (Thermo Fisher Scientific) according to manufacturer instructions. RNA was quality controlled by Qubit fluorometer (Thermo Fisher Scientific) and 2100 Bioanalyzer (Agilent). Samples with RNA integrity number (RIN)-values below 8 were discarded. RNA was transcribed to full-length cDNA using the SMART-Seq2^53^ protocol and RNA-sequencing libraries were generated using the NEBNext^®^ Ultra™ II FS DNA library preparation kit (New England Biolabs) according the manufacturer’s protocol with DNA input below 100 ng. Libraries were indexed using unique i5 and i7 combinations and equimolarly pooled into one multiplex. The multiplex was sequenced over two lanes on a NovaSeq 6000 using the S1 100 cycle paired-end kit generating approximately 35×10^6^ reads per sample.

### Whole genome bisulfite sequencing

For whole genome bisulfite sequencing (WGBS) analysis of dormant and proliferative TC, lungs of 6 mice were pooled into one sample and 200.000 proliferative TC as well as total dormant TC were sorted from 4 pools. Sorted samples were centrifuged and cell pellets were snap frozen on dry ice. Genomic DNA was extracted using the NucleoSpin tissue mini kit for DNA from cells and tissue (Macherey-Nagel). DNA integrity was assessed by TapeStation (Agilent) and samples with DNA integrity number (DIN)-values below 7 were discarded. WGBS libraries were prepared using the xGen™ Methyl-Seq DNA Library Prep Kit (IDT) with partially modified steps in bead clean-up/size selection. Briefly, 200ng genomic DNA was fragmented to 700 – 1000 bp using Covaris ultrasonicator (Covaris, Inc.) and quality checked using TapeStation (Agilent Technologies). Fragmented DNA samples were treated with bisulfite using the EpiTect Bisulfite Kit (Qiagen) following the instructions in the Illumina WGBS for Methylation Analysis Guide (Part # 15021861 Rev. B). After bisulfite conversion adapters were attached to 3`ends of single-stranded DNA fragments which are then extended to generate complementary uracil-free molecules as described in manufactures protocol. The double-stranded DNA fragments were subsequently cleaned up using 1.6x AMPure XP beads (Beckman Coulter) and size selected with a bead ratio of 0.6x and 0.2x, followed by ligation of truncated adapter 2 to uracil-free strand. The adapter-ligated libraries were enriched and indexed using 6 cycles of PCR and purified using magnetic beads according to the protocol. Amplified libraries were quality checked using Qubit fluorometer (Thermo Fisher Scientific) and TapeStation (Agilent). Libraries were pooled equimolarly into one multiplex and sequenced over two lanes on a NovaSeq 6000 using the S1 150 cycle paired-end kit enabling an average genomic coverage of >15.

### Niche-labelling RNA-Sequencing analysis

Raw sequencing data were demultiplexed and FASTQ files were generated using bcl2fastq software (Illumina, v2.20.0.422). FASTQ files were mapped to the GRCm38 mouse reference genome using salmon (v0.7.2)^54^ and count matrices were constructed with the R package tximport (v1.18.0)^55^. Differential gene expression analysis was performed using DESeq2 (v.1.30.1)^56^. Each condition was tested against each condition and differentially expressed genes were used for gene set enrichment analysis (GSEA). GSEA^57,58^ was performed using the R package clusterProfiler (v3.18.1)^59^ or the GSEA java desktop application and the Molecular Signatures Database (MSigDB, v7.4)^32^ provided by the Broad Institute.

For the proliferative niche EC gene panel, DEG were filtered for genes that were specifically upregulated in proliferative niche EC in at least three comparisons (*p* < 0.01 & log2 fold change > 0.5) and were not regulated in non-proliferative niche EC comparisons. For deconvolution of bulk samples aCap and gCap marker genes were defined using the scRNAseq dataset. Expression coefficients (aCap/gCap) of summed marker genes were calculated for each bulk sample using quasibinomal fitting and normalised to PBS injected control samples. Resulting ratios were exponentiated for plotting.

### Single-cell RNA-seq analysis

#### Pre-processing and normalisation

Raw sequencing data were processed as described above. Gene expression was normalised to the mean expression of a housekeeping gene panel (*Actb, Gapdh, Tubb5, Ppia, Ywhaz, B2m, Pgk1, Tbo, Arbp, Gusb, Hprt1*) for each cell, scaled with factor 10.000 and log10 normalised. Normalised count matrices were further analysed using the R package Seurat (v4.0.1)^60,61^. Gene counts per cell, read counts per cell and percentage of mitochondrial transcripts were computed using the respective functions of the Seurat package. For the EC dataset, cells with a percentage of mitochondrial transcripts greater than 5%, along with those with fewer than 1000 genes were excluded. For the TC dataset, only cells with a mitochondrial transcript percentage less than 5% and more than 2500 genes were kept for further analysis.

#### Dimension reduction and clustering

Shared nearest neighbour (SNN)-based clustering and UMAP visualization were performed using the FindClusters and RunUMAP functions within the Seurat package. Each of these were performed on the basis of a principal component analysis, which was performed using the RunPCA function of the Seurat package. For the EC dataset dimensional reduction was performed on 10 principal components (PC) with the resolution parameter set to 0.2 for clustering, whereas for the TC dataset 15 PCs and 0.5 resolution were used. Clusters were annotated using congruent marker expression or according to enrichment for cells derived from a specific FACS gate. Contaminating cells were removed from the dataset based on expression of immune marker genes (*Ptprc, Itgam, Itgax, Adgre1, Cd3e, Cd19, Cd56*) or stromal cell and vessel mural cell marker genes (*Pdgfrb, Des, Myh11, Col1a2, Pdgfra, Cspg4, Pdpn, Acta2*)

#### Cell cycle scoring and differential gene expression

The cell-cycle state was assessed using the gene set and scoring system described previously^62^. Briefly, the S and G_2_M scores were calculated based on a list of 43 S phase-specific and 54 G_2_ or M phase-specific genes. Cells that originated from the dormant FACS gate and had summed scores of less than −1 were tested against cells from the proliferative gate that had score sums greater than 0. Differentially expressed genes were calculated by Wilcoxon rank sum test using FindMarkers-function in Seurat and used for GSEA as described above.

EMT and Wnt gene sets were compiled from MSigDB and gene scores for each cell were calculated using the AddModuleScore-function in Seurat.

#### Trajectory analysis

Trajectory analysis of lung resident TC was performed using the R package Monocle (v.3 alpha)^63,64^. The filtered and normalised TC count matrix was subset from the TC Seurat object. Clustering and dimension reduction was performed using default parameters in Monocle3. The trajectory graph was built by setting cells from the intravascular sorting-gate as starting point. Cells were coloured according to cluster identities as identified in Seurat. Gene expression of EMT and Wnt gene sets was visualised using the plot_cells function.

#### Analysis of TC-EC interactions

Leveraging the biological replicates, TC pseudo-bulks were formed, for which all counts from cells of a specific FACS gate and biological replicate were summed and differentially expressed genes (DEG) were computed for the experimental timepoints using DESeq2^56^. DEG were filtered against CellPhoneDB^27^ database to retrieve putative ligands and receptors. CellPhoneDB ligands and receptors were further filtered for expression of interaction partners in the day 1.5 EC dataset (for intravascular versus extravascular comparison) or for expression in the day 3.5 EC dataset (for dormant versus proliferative comparison), respectively. Log2 fold changes of TC-expressed ligands or receptors were plotted against each. Receptors or ligands with upregulation in extravasated and dormant TC were considered trajectory defining, as well as receptors or ligands upregulated in intravascular and proliferative TC.

#### Analysis of publicly available human CTC datasets

Normalised and filtered count matrices were downloaded from provided source data^30^ or the gene expression omnibus (GEO) under the accession code GSE109761^31^. Dimension reduction and visualization were performed in Seurat as described above using default parameters. Gene scores of human orthologues of the EMT and Wnt gene lists were computed as described above.

### WGBS data analysis

Raw sequencing data were demultiplexed and FASTQ files were generated using bcl2fastq software (Illumina, v2.20.0.422). FASTQ files were trimmed using Trimgalore (v0.6.6)^65^ and mapped to the GRCm39 mouse reference genome using Bismark (v0.22.3)^66^. Forward and reverse strands were collapsed and methylation sites were called in Bismark. Differentially methylated regions were determined with the R package bsseq using default parameters (v1.26.0)^67^. For this, biological replicates were summed and methylation fractions for annotated genomic regions between the two conditions were compared. Regulatory elements, promoters and gene bodies were annotated with annotation sheets downloaded from Ensembl database (release 105)^68^.

### Real time quantitative PCR

Total RNA of cell cultured tumour cells was isolated using the GenElute Mammalian Total RNA Purification Kit (Merck) according to the manufacturer’s instructions. 1000 ng of RNA was reverse transcribed using the QuantiTect Reverse Transcription Kit (Qiagen) according to the manufacturer’s protocol. Gene expression analysis was performed by quantitative PCR using TaqMan reactions (Thermo Fisher Scientific) and Lightcycler 480 (Roche). Gene expression levels were assessed using the C_t_-method and normalised to the expression of *Actb*, resulting in ΔC_t_-values. Relative gene expression was assessed by normalizing ΔC_t_-values of individual samples to the average control ΔC_t_-value, resulting in ΔΔC_t_-values. Relative fold changes to control were then calculated as 2^-ΔΔCt^.

### Histology

Zinc-fixed primary tumours were paraffin embedded and cut into 7 µm sections. Sections were deparaffinized and rehydrated and antigen retrieval was performed by incubation with Proteinase K (20 µg/ml, Gerbu Biotechnik) for 5 min at 37°C. Tissues were blocked in 10% ready-to-use goat serum (Zymed) for 1 hour at room temperature, followed by overnight incubation with rat anti-CD31 (1:100, BD Bioscience) and rabbit anti-Desmin (1:100, Abcam) diluted in blocking buffer at 4 °C. After three washes in TBS-T, slides were stained with anti-rat Alexa647 and anti-rabbit Alexa546 antibody at room temperature for 1 hr. Cell nuclei were counterstained with 1:2000 Hoechst 33342 (Sigma-Aldrich) and sections were mounted with DAKO mounting medium (Agilent). Images were acquired as whole-area tile scans using an Axio Scan.Z1 slide scanner (Zeiss). Image analysis was performed using Fiji software (ImageJ, 1.53q). After region-of-interest (ROI) selection, CD31, Desmin and DAPI channels were binarized using thresholding. For vessel area, percentage of CD31+ area within the ROI was calculated. For vessel coverage, CD31 channel was masked and Desmin overlap with CD31 was calculated. CD31+/Desmin+ double-positive vessel were considered covered and coverage was calculated as the ratio of covered/uncovered vessels.

### Statistical Analysis

Statistical analysis was performed using GraphPad Prism (v6) and R (v4.0.5).

